# A human sensory neuron model for varicella-zoster virus latency and reactivation *in vitro*

**DOI:** 10.64898/2025.12.12.693952

**Authors:** Alexander C. Havelaar, Joseph S. Flot, Saori Fukuda, Anja W.M. de Jong, R.I. Koning, Amber Schotting, Sem van ‘t Geloof, Tomohiko Sadaoka, Georges M.G.M. Verjans, Paul R. Kinchington, Werner J.D. Ouwendijk

**Author notes:** These authors contributed equally (Alexander C. Havelaar, Joseph S. Flot).

## Abstract

Varicella-zoster virus (VZV) is a ubiquitous human neurotropic alphaherpesvirus that establishes lifelong latency in sensory ganglionic neurons. Subsequent viral reactivation causes herpes zoster, a morbid disease often complicated by neuropathic pain. Mechanisms underlying VZV latency and reactivation are not understood, mostly due to the lack of permissive animal models and challenges of current *in vitro* latency modelling. Here, we evaluated HD10.6 cells, a simplified and easily expandable human sensory neuron line to model VZV latency and reactivation. Mature HD10.6 (mHD10.6) differentiated neurons supported productive VZV infection, viral DNA replication, production of infectious progeny, and viral spread in cultures. VZV infection was associated with limited cytopathic effects and ultrastructural changes. Infecting mHD10.6 neurons with cell-free VZV in the presence of antivirals resulted in a quiescent-persistent state, characterized by persistent VZV genomes with restricted VZV gene expression and absence of infectious virus. Importantly, VZV could be reactivated by treatment with capsaicin, as evidenced by increased lytic viral transcription and virus spread. In conclusion, this study establishes human HD10.6 neurons as a novel and scalable *in vitro* model for studying VZV latency and reactivation to identify virus and host factors governing latency that may serve as therapeutic targets to restrict VZV reactivation.

**Importance:** Most individuals carry latent varicella-zoster virus (VZV) in their dorsal root ganglia (DRG), which can reactivate to cause shingles and chronic pain. The mechanisms by which VZV establishes latency and triggers of reactivation are incompletely understood. Current platforms for the study of VZV latency do not easily support functional experiments (ganglia) or are difficult to expand and complicated by mixed neuronal populations (stem cell-derived neurons). Here, we demonstrate that matured HD10.6 (mHD10.6) cells derived from immortalized human DRG-derived neurons provide a clonal, scalable, and easy-to-expand platform for studying lytic, latent, and reactivated VZV infection in sensory neurons. We propose that the HD10.6 platform could provide the basis to conduct studies on the viral latent state that have hitherto not been possible.

## Introduction

Varicella-zoster virus (VZV) is an ubiquitous human neurotropic alphaherpesvirus^1^. Primary VZV infection causes varicella (chickenpox), followed by the establishment of a lifelong latent infection in sensory neurons of the dorsal root ganglia (DRG) and trigeminal ganglia (TG)^2–4^. Later in life, the virus reactivates in about one-third of infected individuals to cause herpes zoster (shingles), which is often morbid, complicated, and frequently followed by difficult-to-treat chronic pain states termed post-herpetic neuralgia (PHN)^5^. The molecular mechanisms underlying VZV latency and reactivation are incompletely understood, largely due to a lack of tractable VZV-permissive animal models and limitations/challenges of current *in vitro* systems to study these processes.

VZV is an exclusively human-restricted pathogen, and therefore much of our current understanding of viral latency has derived from studying post-mortem human ganglia. Latent VZV DNA resides within the nuclei of an average of 2-5% of sensory neurons per ganglion, primarily in an episomal form^6–10^. The latent VZV genome is transcriptionally repressed, likely due to the deposition of repressive chromatin on viral DNA^11^. As such, expression of the approximately 136 lytic viral RNAs and the proteins they encode is blocked. An exception is the VZV latency-associated transcript (VLT) and two isoforms of VLT-ORF63 transcripts that have been identified in human TG^11–14^. While studies on human TG obtained at short post-mortem intervals capture the latent VZV state *in vivo*, their use is hitherto limited to observational studies. Therefore, additional model systems are required to study the roles of VLT and VLT-ORF63 RNAs and the drivers of reactivation.

Various neuronal *in vitro* models of human origin have thus been developed to pursue functional latency/reactivation studies^15,16^. These include cultured human neurons developed from human induced pluripotent (hiPSC) or human embryonic (hESC) stem cells^17–19^, commercially available hiPSC-derived sensory neuron progenitors (HSNP)^20^, and the neuroblastoma cell line SH-SY5Y^21–23^. SH-SY5Y are immortalized neuroblastoma cells that can be differentiated into dopaminergic central nervous system (CNS) neuron-like cells, which support lytic VZV infection and possibly a reactivatable latent-like state^22,23^. hiPSC-, hESC-and HSNP-derived neurons can host model latent states after lytic suppression using antivirals or following axonal infection in microfluidic chambered neurons. Reactivation can be experimentally induced by various strategies, including interruption of nerve growth factor (NGF) signaling, agents that induce chromatin remodeling, or by VLT-ORF63 overexpression^17–19^. Some of these systems generate multiple neuron subtypes, with many resembling CNS-like neurons^17–19^. While HSN more closely resemble human DRG neurons, these and other stem cell-derived neuron models are expensive, challenging to work with, and difficult to scale up to large-scale neuron cultures. There remains a need for a simpler platform for a deeper research probing of latency.

To address the limitations of existing *in vitro* models, we explored the use of the immortalized human DRG-derived clonal cell line HD10.6^24^. HD10.6 cells are easily expandable committed DRG neuron precursors that contain a tetracycline-regulatable (Tet-off) v-*myc* oncogene. In the presence of doxycycline and specific neuronal growth factors, they differentiate into mature sensory DRG neurons, which are capable of firing action potentials. They are sensitive to capsaicin, a hallmark of nociceptive neurons^25–27^. Previously, it was shown that mature HD10.6 cells support lytic herpes simplex virus 1 (HSV-1) infection and model latent states, from which HSV-1 could be stimulated to reactivate^25,28^. Here, we aim to develop these cells into a model for VZV latency and reactivation.

## Results

### HD10.6 cells differentiate into human mature sensory DRG neurons

We performed short tandem repeat (STR) profiling to confirm the identity of the HD10.6 cells and expand the currently known STR profile (**Supplementary Table 1**). To validate that we could differentiate immature HD10.6 (iHD10.6) cells into human mature sensory neurons with similar kinetics as previously reported, HD10.6 cells were seeded on Matrigel and poly-D-lysine coated cell culture plates and cultured in maturation medium for 10-21 days (**Fig. 1A**). Consistent with Thellman *et al*, the addition of doxycycline and neurotrophins rapidly induced visible signs of differentiation, including the formation of axon-like projections, which started to form networks at 6 days of differentiation and were clearly apparent at day 10 and 21 of differentiation (**Fig. 1B**)^26^. Differentiated HD10.6 cells (mature HD10.6; mHD10.6) at day 10 expressed the neuron marker β-III-tubulin (*TUBB3*), the peripheral sensory neuron marker peripherin (*PRPH*), the sensory neuron marker *RET*, and neurotrophin receptor TrkB, as shown by immunofluorescent staining (**Fig 1C**) and/or RT-qPCR (**Fig. 1D**). To validate that iHD10.6 cells differentiate into a pure population of mature sensory neurons, but not non-neuronal cells, we performed RT-qPCR for neuronal markers (*NTRK2*, *PRPH*, *NEFH*), satellite glial cells (*ITGAM*), Schwann cells (*MOBP*) and fibroblasts (*COL1A2*) on mHD10.6 neurons and human cadaveric TG obtained with a short post-mortem interval. Whereas neuronal gene expression was abundant in mHD10.6 cells, the expression of glia- and fibroblast-specific genes was found only in the human cadaver TG sample and was nearly absent in the HD10.6 cultures (**Fig. 1E**). Additionally, no cells expressing microglia marker IBA1 or Schwann cell marker GFAP were observed in immature and mature HD10.6 cultures by immunofluorescent staining (data not shown). Thus, the human fetal DRG-derived HD10.6 cells provide an immortalized, clonal, scalable, and pure sensory neuron model to study VZV infection.

**Figure 1.**
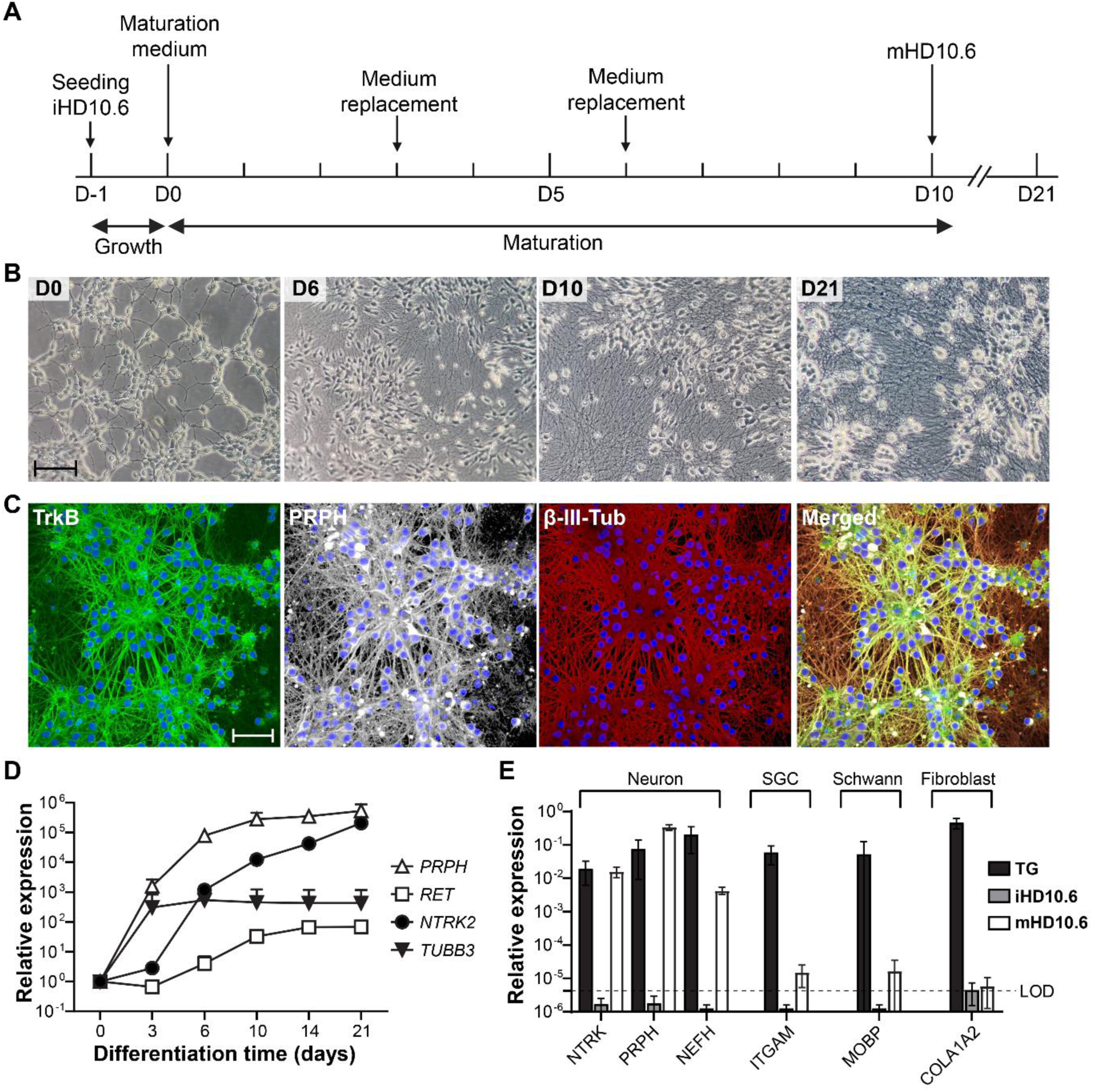
Differentiation of H10.6 cells into mature peripheral sensory neurons. (**A**) Timeline of the maturation process from immature HD10.6 cells into mature HD10.6 neurons. (**B**) Representative bright-field microscopy images of HD10.6 cells between 0 and 21 days post-maturation. (**C**) Representative confocal microscopy images of mHD10.6 (10 days post-maturation) stained for TrkB (green), peripherin (white), and beta-3-tubulin (red) and counterstained with Hoechst (blue) upon differentiation. Scale bar = 100 µm (**D**) Graph showing relative transcription levels of *PRPH* (peripherin), *RET*, *NTRK2* (TrkB), and *TUBB3* (beta-3-tubulin), normalized to β-actin transcription levels, between 0 and 21 days of differentiation, determined by RT-qPCR. Data shown are mean ± SD of three independent experiments. (**E**) Graph showing relative expression of markers for sensory neurons (*NTRK2*, *PRPH*, *NEFH*), satellite glia cells (SGC; *ITGAM*), Schwann cells (*MOBP*), and fibroblasts (*COL1A2*) in human TG, iHD10.6, and mHD10.6 cells (mHD10.6 10 days post-maturation), normalized to β-actin expression measured by RT-qPCR. Data are shown for n=6 TG, n=1 iHD10.6, and n=2 mHD10.6 cultures in triplicate. LOD, limit of detection. Bars and error bars indicate mean ± SD.

### Generation of recombinant pOka-derived VZV-GFP viruses

To facilitate visualization of VZV spread upon infection of mature HD10.6 (mHD10.6) neurons, we generated a recombinant VZV strain expressing a GFP reporter fused in-frame to the ORF49 gene. The ORF49-T2A-GFP cassette was inserted into the ORF49 locus of the pOka BAC strain by recA-mediated allelic exchange (**Fig. 2A**), as described^29^. ORF49 is one of the most abundantly transcribed VZV genes, is expressed with *true-late* kinetics, and encodes for a non-essential tegument protein that functions in virion maturation^13,30–32^. To ensure that the resulting VZV-ORF49-T2A-GFP (VZV-49GFP) was not attenuated for lytic VZV infection, we infected highly susceptible human retinal pigment epithelial (ARPE-19) cells and analyzed virus replication by multistep growth curve analysis. VZV infection in ARPE-19 cells resulted in bright GFP expression in culture (**Fig. 2B**). Importantly, viral DNA levels and infectious cell-associated virus titers were similar between VZV-49GFP and its parental wild-type VZV rpOka virus, which indicates that VZV-49GFP is not attenuated compared to wild-type virus (**Fig. 2C-D**). A second GFP reporter virus was used in some later studies, which contained a GFP-luciferase cassette under the ORF57 protein and promoter (VZV-57LZ-GFP). This virus was derived without BAC recombination, and its derivation will shortly be detailed in a separate manuscript (Flot and Kinchington, in Preparation).

**Figure 2.**
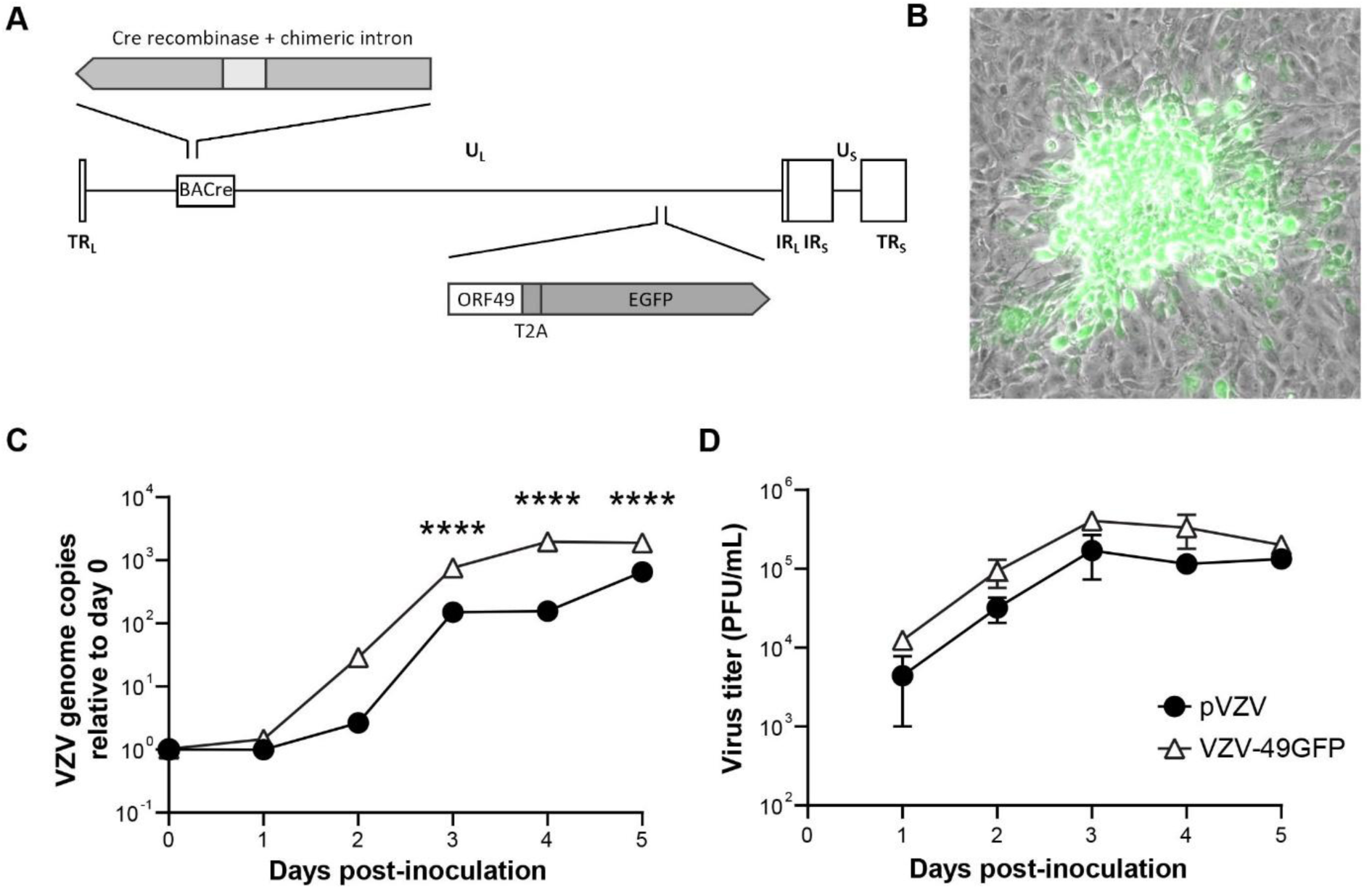
Generation and replication kinetics of VZV-49GFP. (**A**) VZV-49GFP was generated by recA-mediated allelic exchange to insert a cassette consisting of the ORF49 gene, a T2A ribosome skipping motif, and EGFP reporter into the native ORF49 locus within the VZV rpOka BACre genome. (**B**) Representative merged fluorescence and brightfield microscopy images showing GFP expression by VZV-49GFP-infected ARPE-19 cells. (**C - D**) Graphs showing growth kinetics of parental VZV BAC (pVZV) and VZV-49GFP, as measured by viral DNA qPCR (**C**) and infectious virus titers (**D**) within infected cells. Data indicate n=2 biological replicates performed in technical triplicates. Symbols and error bars indicate mean ± SD.

### Mature HD10.6 neurons support productive VZV infection

To determine whether mHD10.6 neurons support productive (lytic) VZV infection, HD10.6 cells were matured for 10 days and infected with cell-free VZV-49GFP at a low multiplicity of infection (MOI; 1000 plaque-forming units (PFU) per well, determined on ARPE-19, corresponding to MOI ± 0.01). This resulted in robust expression of VZV immediate early (*IE*; ORF4 and ORF63), early (*E*; ORF29), and early-late (*EL*; ORF68 and VLT) genes at 4 days post-infection (dpi) (**Supplementary Fig. 1**) and increased levels of VZV DNA over time (**Fig. 3A**), indicative of viral DNA replication. Newly produced VZV progeny capable of infecting ARPE-19 cells remained associated with infected neurons (**Fig. 3B**). Consistent with cell-to-cell spread of VZV to neighboring uninfected neurons, the size, but not the number, of GFP-positive infectious foci increased over time (**Fig. 3C**). Interestingly, the GFP-positive VZV-infected mHD10.6 neurons appeared visually similar to mock-infected neurons, with little apparent cytopathic effect (CPE) or cell death until at least 10 dpi (**Fig. 3D)**. This suggests VZV infection causes minimal cytopathic effects in HD10.6 cells despite the production of infectious virus.

**Figure 3.**
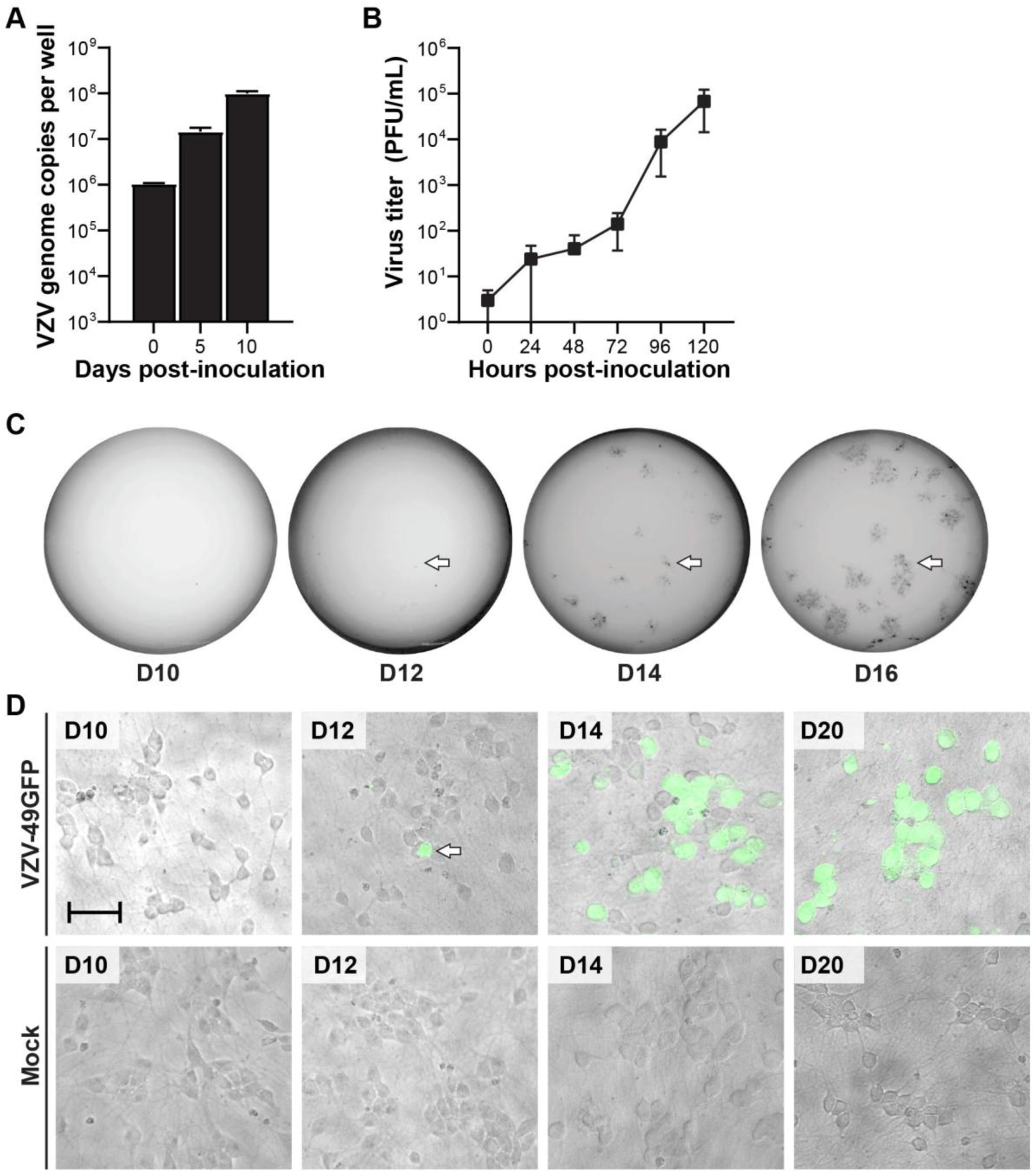
mHD10.6 neurons support productive VZV infection. (**A**) VZV DNA qPCR on DNA extracted from VZV-49GFP-infected mHD10.6 neurons (n=3 biological replicates in technical triplicates). Bars and error bars indicate mean ± SD. (**B**) Cell-associated infectious virus titers as determined by titration of VZV-49GFP-infected mHD10.6 neurons on ARPE-19 cells (n=3 biological replicates in technical triplicates). Symbols and error bars indicate mean ± SD. (**C**) Images showing GFP expression in VZV-49GFP-infected mHD10.6 neurons at multiple times after infection, as measured by the Typhoon fluorescence imaging system (representative images shown for n≥3 biological replicates in technical triplicates). Arrow indicates same GFP-positive focus. (**D**) Representative merged fluorescence and brightfield microscopy images of uninfected and VZV-49GFP-infected mHD10.6 neurons (representative images shown for n≥3 biological replicates in technical triplicates). Scale bar 100 µm.

### VZV induces ultrastructural changes and forms abundant virions in infected mHD10.6 neurons

To more deeply investigate the morphological changes induced by VZV infection, we analyzed mock- and VZV-49GFP-infected mHD10.6 neurons at 7 dpi at the ultrastructural level by transmission electron microscopy (TEM). Viral capsids (**Fig. 4A-C)** and particles (**Fig. 4D-F)** were readily observed in the nucleus, cytoplasm, and on the plasma membrane of VZV-infected HD10.6 neurons. Different stages of capsid assembly and maturation were captured, including empty capsids (**Fig. 4A**), capsids still apparently containing internal structures that may be scaffold proteins (**Fig. 4B**) and complete mature capsids with apparent packaged VZV DNA (**Fig. 4C**). Abundant VZV particles were observed on the cell surface of neuronal cell bodies and axons, which were composed of defective particles lacking a nucleocapsid (**Fig. 4D**), virions that incorporated defective capsids lacking dense cores (**Fig. 4E**) and complete VZV virions containing a capsid with a central dense core, likely of DNA (**Fig. 4F**). Consistent with studies on VZV infection in stem-cell derived neurons^33^, we observed that about 10% (n=±1100 virions analyzed) of surface virions were complete.

**Figure 4.**
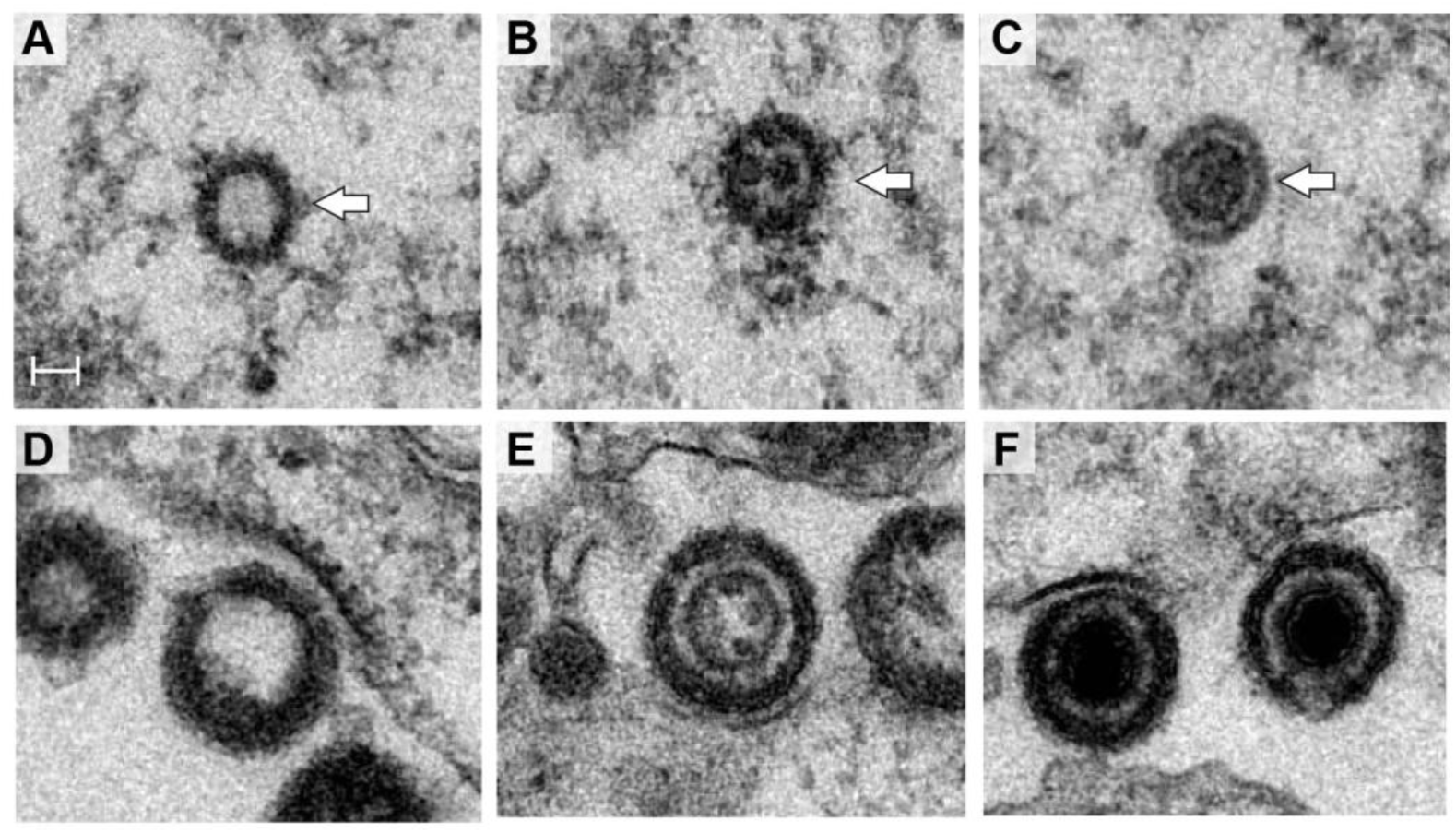
Transmission electron microscopy (TEM) analysis of VZV capsids and particles in VZV-infected mHD10.6 neurons. (**A-C**) Representative TEM images showing empty capsid (**A**), capsid with scaffolding proteins (**B**), and nucleocapsid (**C**) in the nucleus at 7 days post-infection (arrows). (**D-F**) Representative TEM images showing empty virions (**D**), a virion that included a scaffold capsid (**E**), and complete VZV virions (**F**) on the plasma membrane. Scale bar 50 nm.

In line with the minimal morphological alterations observed in macroscopic images of VZV-infected HD10.6 neurons, the majority of infected cells showed only mild morphological changes and little cellular disruption and disorganization using electron microscopy (**Fig. 5A-H**). Accumulation of electron-dense material in the Golgi apparatus and dilation of the endoplasmic reticulum (ER) were apparent (**Fig. 5G**), in contrast to uninfected mHD10.6 (**Fig. 5C)**, but we saw no nuclear deformation **(Fig. 5F)** similar to uninfected controls (**Fig. 5B**). Occasionally, more advanced structural changes were observed (**Fig. 5I)**, including nuclear condensation (**Fig. 5J)** and marked expansion of the ER and Golgi, giving the cytoplasm a vacuolated appearance (**Fig. 5K**). Infected neurons showed abundant particles at the apparent outer surface **(Fig. 5E)**. These observations indicate efficient assembly and egress of VZV in mHD10.6 neurons, with relatively limited cellular damage.

**Figure 5.**
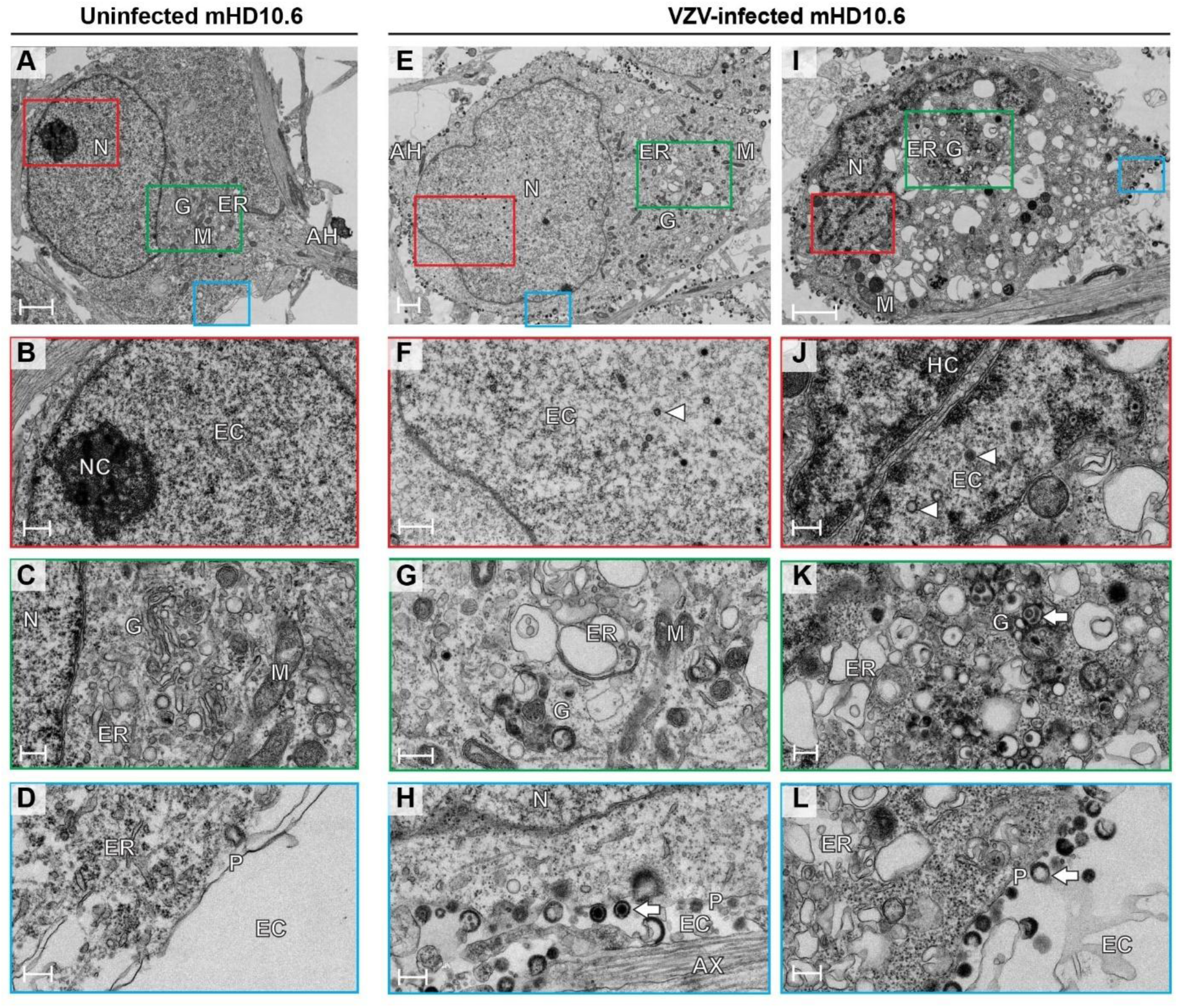
Transmission electron microscopy (TEM) analysis of ultrastructural changes in VZV-infected mHD10.6 neurons. Representative TEM images of uninfected (**A-D**) and VZV-infected mHD10.6 neurons at 7 days post-infection (**E-L**). Nucleus (N), Golgi apparatus (G), endoplasmic reticulum (ER), mitochondria (M), and the axon hillock (AH), axon (AX), plasma membrane (P), nucleolus (NC), and euchromatin (EC) are, where applicable, indicated. VZV capsids (**F, J**) are indicated by arrowheads, and particles (**K, H, L**) are indicated by arrows. Scale bar: 1 µm (A, E, I), 500 nm (B, F) or 250 nm (C, D, G, H, J, K, L).

### Mature HD10.6 neurons support a quiescent-persistent VZV infection

To determine if HD10.6 matured neurons can host a non-productive persistent (and potentially latency-like) VZV infection, we infected mature HD10.6 neurons with cell-free VZV-49GFP at 1000 PFU/well (MOI ± 0.01) in the presence of antivirals for 5 days, followed by 5 days without antiviral (**Fig. 6A**). In addition to the nucleoside analog acyclovir (ACV, 300 µM), which is commonly used to establish quiescent VZV and HSV-1 infections *in vitro*^34,35^, we also tested the nucleoside analog brivudine (BVDU at 2 µM), which has high potency against VZV^36–38^, and the helicase-primase inhibitor amenamevir (AMNV, 1 µM)^39–43^. Comparable results were obtained for ACV, BVDU, and AMNV in that all inhibited VZV infection, with AMNV being slightly less effective at silencing gene transcription compared to ACV and BVDU at the concentrations tested (**Fig. 6** and **Supplementary Fig. 2**). Intriguingly, all inhibited the expression of GFP during antiviral treatment, but occasional GFP-positive cells appeared upon removal of antivirals (**Fig. 6C**). However, such individual GFP-positive cells did not show spread to other cells over time, suggesting that full lytic replication to virus production was blocked. Indeed, to confirm that no infectious virus was produced, we co-cultured lytic or antiviral-treated/removed latently-infected mHD10.6 neurons with ARPE-19 cells after 5 days of infection. In contrast to lytic VZV infections, we could never recover infectious virus from quiescently-infected mHD10.6 cultures and supernatant during antiviral treatment or after the period of removal of antivirals (**Fig. 6B**). To investigate if the VZV genomes persisted in such cultures and were transcriptionally repressed, we performed RT-qPCR to quantify viral transcripts and VZV genomic qPCR to measure viral DNA load in lytic and latent VZV-infected mHD10.6 cultures. Compared to lytic infection, gene expression of *IE* genes ORF4 and ORF63, the *E* gene ORF29, and *EL* gene ORF68 were all strongly repressed, but not absent (**Fig. 6D** and **Supplementary Fig. 2**). Surprisingly, expression of *VLT*, a hallmark of VZV latency in human latently infected TG, was similarly reduced. VZV DNA loads, however, remained stable over time, strongly indicating that no viral DNA replication occurred and that VZV DNA persisted within infected mHD10.6 cells (**Fig. 6E** and **Supplementary Fig. 2B**).

**Figure 6.**
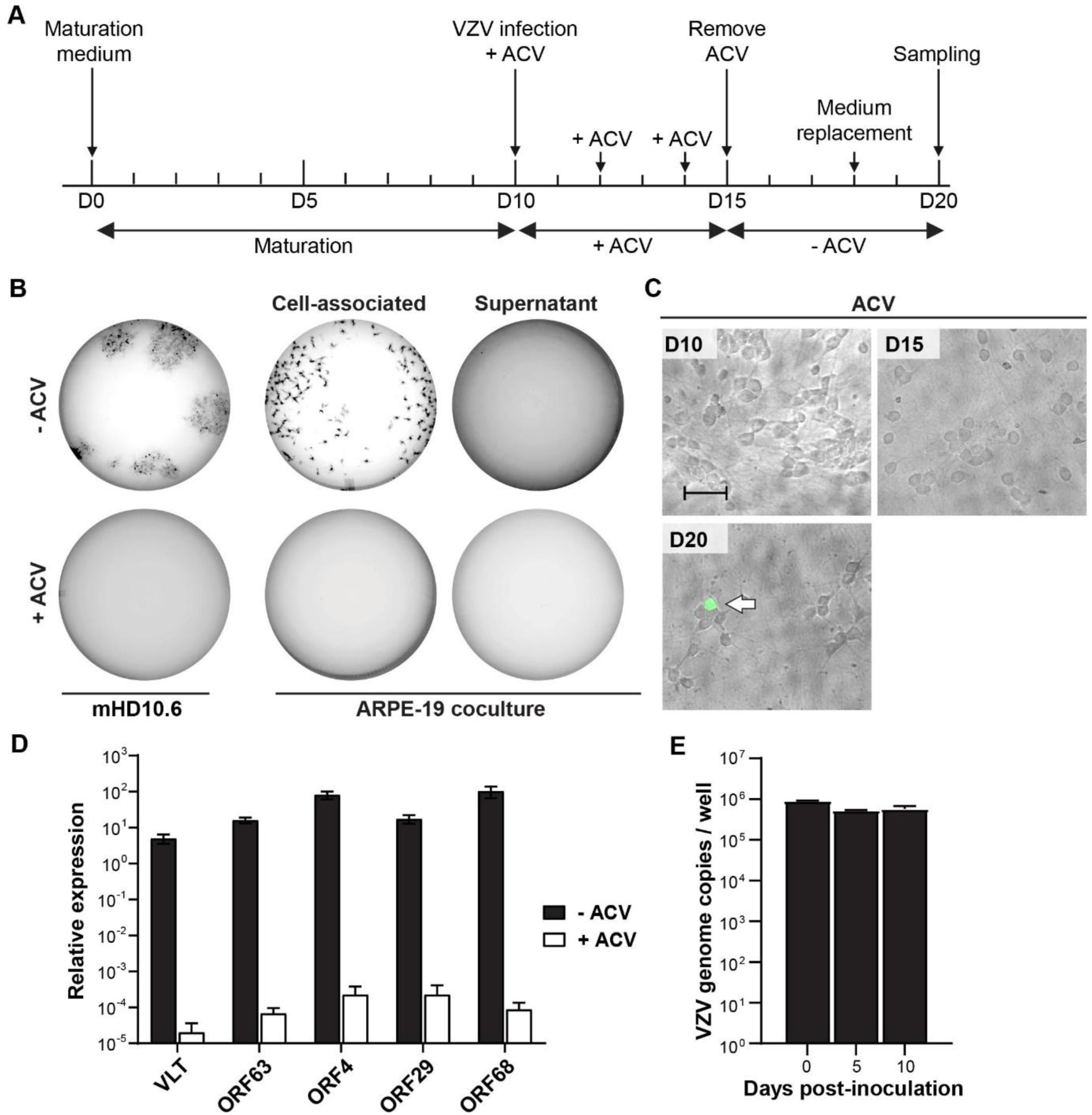
Characterization of quiescent-persistent VZV infection of mHD10.6 neurons at 10 dpi. (**A**) Timeline showing the outline of the experiment. (**B**) Images of GFP expression in VZV-49GFP-infected mHD10.6 neurons, and ARPE-19 cells co-cultured with neurons that were infected in the presence or absence of ACV, measured by the Typhoon fluorescence imaging system (n=3 biological replicates performed in triplicate). (**C**) Representative merged fluorescence and brightfield microscopy images showing that no GFP-positive neurons were present during ACV treatment (days 0 and 5), but occasional single GFP-positive cells (arrow) appeared upon ACV removal (day 10) (n = 3 biological replicates performed in triplicate). (**D**) RT-qPCR analysis of the indicated VZV transcripts in lytic (10 dpi) and latent (10 dpi) mHD10.6 neurons. Data are expressed as relative transcript level compared with housekeeping gene β-actin using the 2^-ΔCt^ method (n=6 biological replicates performed in triplicate). (**E**) VZV DNA qPCR on DNA extracted from latent VZV-49GFP-infected mHD10.6 neurons (n=3 biological replicates performed in triplicate). Bars and error bars indicate mean ± SD.

We hypothesized that a longer antiviral treatment may promote the establishment of a more latency-like repressed state of the viral genome. Therefore, we infected mHD10.6 neurons in the presence of 300 µM ACV and analyzed cultures after 1 or 10 days of ACV treatment (day 11; day 20) and 10 days after ACV removal (day 30) (**Fig. 7A**) for the presence of viral transcripts and DNA by (RT-)qPCR. While ACV treatment significantly reduced VZV transcript levels at 1 dpi compared to no ACV treatment, the 10-day ACV treatment (day 20) showed much reduced transcripts that did not return after 10 days following ACV removal (day 30) (**Fig. 7B-C**). Interestingly, VZV transcripts originating from the VLT and ORF63 loci appeared to decline to a lesser degree than lytic genes ORF4, ORF29 and ORF68 (**Fig. 7B** and **Supplementary Fig. 3**). By contrast, lytic VZV infection, was associated with increased viral RNA levels over time (**Fig. 7B)**, and while DNA load was similar between lytic and ACV-treated cultures at day 1, they remained stable during ACV treatment, and showed a mild decrease at day 20 compared to day 1 (**Fig. 7C**). Again, we did not observe any GFP-positive cells in the presence of antivirals, but upon removal of the antivirals between 1–5 GFP-positive cells per well appeared. These GFP-positive cells remained GFP-positive, but did not spread throughout the cultures. Combined, these data indicate that VZV infection of mHD10.6 cells in the presence of antivirals results in a quiescent-persistent state, in which there is persistence of viral DNA in conjunction with repressed viral gene expression and no replication or new infectious virus.

**Figure 7.**
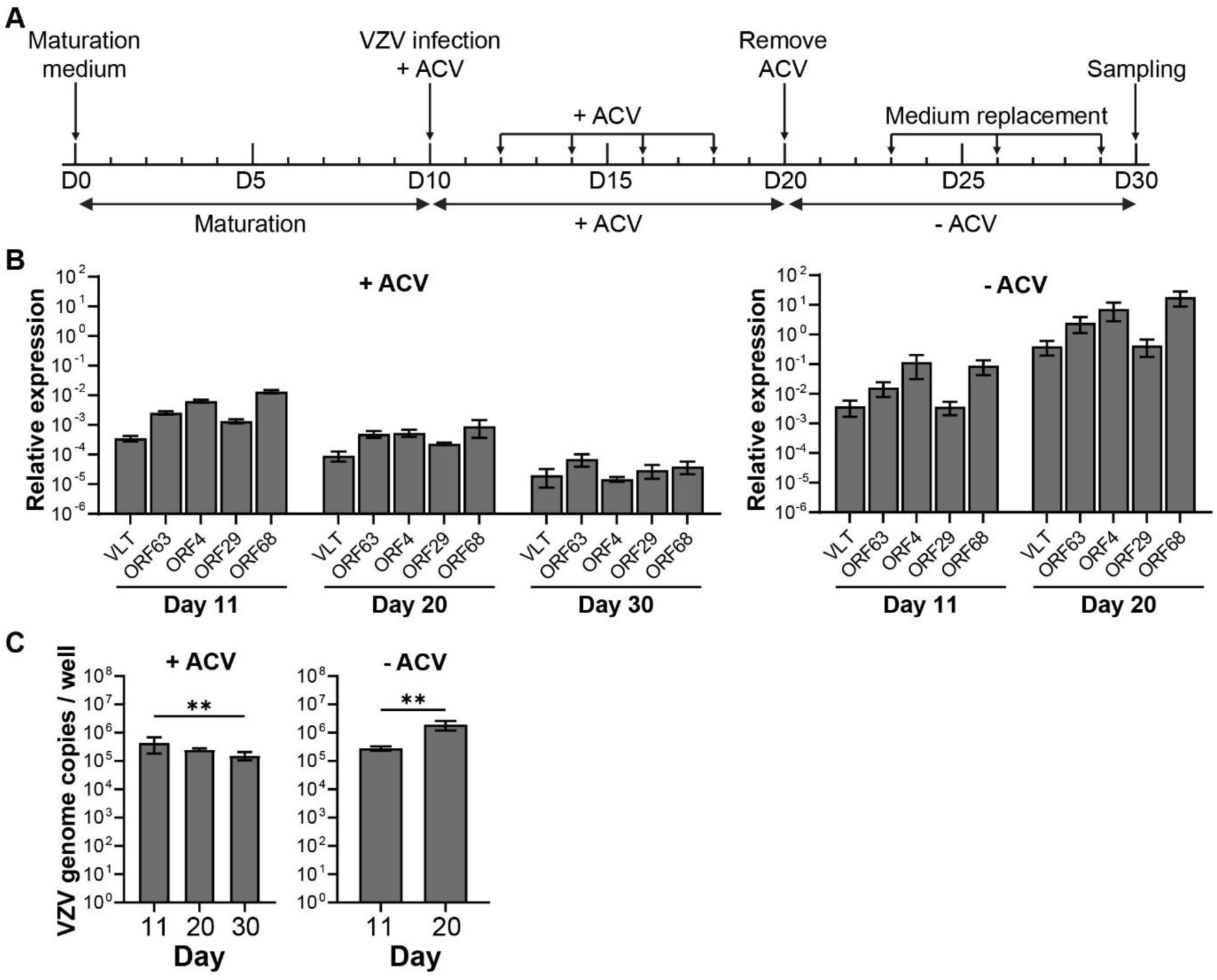
Characterization of quiescent-persistent VZV infection of mHD10.6 neurons at 20 dpi. (**A**) Timeline showing the outline of the experiment. (**B**) RT-qPCR analysis of the indicated VZV transcripts in lytic and latent mHD10.6 neurons. Data are expressed as relative transcript level compared with housekeeping gene β-actin using the 2^-ΔCt^ method (n=3 biological replicates performed in triplicate). (**C**) VZV DNA qPCR on DNA extracted from lytic and latent VZV-49GFP-infected mHD10.6 neurons. Bars and error bars indicate mean ± SD.

### Mature HD10.6 cells support viral reactivation of VZV

We have established that mHD10.6 neurons can host silenced VZV genomes induced by antivirals. Accordingly, we next determined the ability of quiescent-persistent VZV-infected mHD10.6 neurons to support virus reactivation. Following a 24-hour pre-treatment in media containing a combination of 300 µM ACV, 2 µM BVDU, and 1 µM AMNV to suppress lytic infection, 24-well cultures of mature HD10.6 were infected with VZV expressing GFP (VZV-57LZ-GFP). Six wells were not treated with antivirals and showed rapid lytic growth and virus spread as detected by imaging of GFP in the cultures. The antiviral regimen was maintained on cultured infected mHD10.6 neurons for 10 days, followed by a withdrawal period in which antivirals were withdrawn, and infected cultures were maintained in differentiation media. Infected neuron cultures were monitored over time for possible breakthrough infections, indicating spontaneous reactivation events **(Fig. 8A)**. We then applied several reactivation stimuli for 48 hours with reapplication after 24 hours, except for inducers with longer stimulation periods **(Table 1)**. Cultures were monitored with whole-well live cell microscopy scans to track the appearance of new GFP-positive foci and spreading infection **(Fig. 8B)**. Over multiple experiments, a fraction of infected mHD10.6 wells treated with capsaicin showed reactivation events, while other reactivation treatments did not result in the appearance of reproducible growing lytic infection events. No cultures left unstimulated showed a sporadic reactivation event. (RT-)qPCR analyses of the reactivated capsaicin treated cultures, and comparison to lytic and latent cultures revealed an increase in the number of viral genome copies, and higher levels of all viral transcripts in capsaicin-reactivated cultures over latently infected, GFP-negative cultures **(Fig. 8C).** Again, we noted detectable levels of viral transcripts in the uninduced cultures, suggesting that some genomes might be partially released from repression without the expression of GFP and virus production. Consistent with a latent state, the HD10.6 neurons also retained detectable VZV genomic DNA, confirming that the cells continued to harbor latent viral genomes (**Fig. 8D**). Taken together, we have demonstrated that HD10.6 matured neurons can hold a stable population of VZV genomes following suppression of lytic events, and treatment with select stimuli can induce reactivation events in the cultures, albeit with low efficiency.

**Figure 8.**
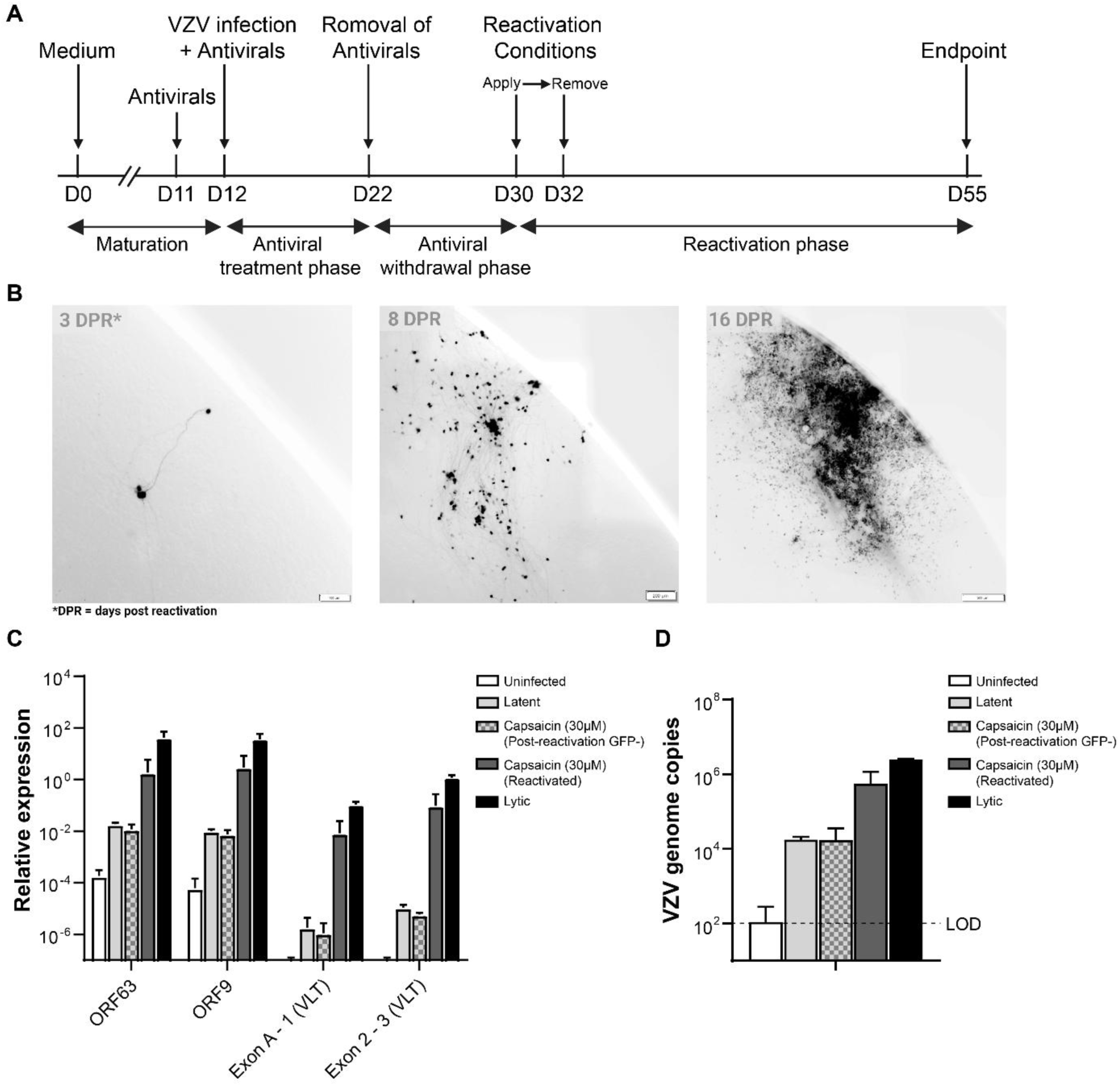
Characterization of mHD10.6 VZV reactivation using capsaicin. (**A**) Timeline showing the outline of one experiment. (**B**) Progression of a reactivated infection originating from a reactivation in a single neuron over the course of 3 - 16 days post reactivation is shown. (**C**) RT-qPCR analysis of the indicated VZV transcripts in lytic, latent, and reactivated mHD10.6 neurons. Data are expressed as relative transcript level compared with housekeeping gene β-actin using the 2-ΔCt method (n=3 to 8 replicates). (**D**) VZV DNA qPCR on DNA extracted from lytic, latent and reactivated VZV-57LZ-GFP-infected mHD10.6 neurons. LOD, limit of detection. Bars and error bars indicate mean ± SD.

**Table 1.** Reactivation stimuli tested on quiescent-persistent VZV-infected mHD10.6 neurons.

| Treatment (concentration) | Exposure time (hours) | Target | Observed reactivations / number of experiments |
| --- | --- | --- | --- |
| LY294002 (20 µm) | 48 | Phosphatidylinositol 3-kinase (PI3-K) inhibitor | 0/3 |
| NGF depletion | 48 | TrkA receptor tyrosine kinase signaling | 0/3 |
| Sodium Butyrate (5 mM) | 168 | Histone deacetylase inhibitor | 0/3 |
| JQ-1 | 48 | BET bromodomain inhibitor | 1/2 |
| Forskolin (60 µM) | 48 | Adenylate cyclase | 0/4 |
| Doxycycline removal | 168 | v-Myc induction | 0/1 |
| ADU-S100 (10 µg/mL) | 48 | Stimulator of interferon genes (STING) activator | 0/2 |
| Capsaicin (30 µM) | 48 | TRPV1 agonist | 2/2 |
| 12-O-Tetradecanoyl-phorbol-13-acetate (TPA) | 168 | Protein Kinase C (PKC) activator | 0/1 |
| Heat Shock (43 °C) | 3 | Heat shock proteins (HSPs) | 0/3 |
| VZV superinfection (MOI 0.1) | 240 | Supply of viral trans-activators | 1/1 |

## Discussion

Probing VZV latency and reactivation has historically been challenging due to the virus’s strict human-specific tropism. While stem cell-derived neuronal models significantly advanced our understanding of some of these processes^14,16–18,20^, they are limited by high cost, limited scalability, and/or lack of peripheral sensory neurons in a mixed population. To complement these models and overcome some of their limitations, we established a novel human peripheral sensory neuron platform of VZV infection, quiescent-persistence, and reactivation based on the immortalized human DRG-derived HD10.6 line.

Mature HD10.6 neurons clearly support lytic (productive) VZV infection, whether it be in conditions without antiviral suppression, or following a period of antiviral treatment followed by stimulation conditions. We confirmed that HD10.6 cells can be matured into peripheral sensory neurons that form elaborate axonal networks. The capsaicin receptor (encoded by the *TRPV1* gene) is expressed by nociceptive sensory neurons, which are small to medium-sized neurons that transmit pain and temperature sensation, and combined comprise the majority of DRG and TG neurons^24,44,45^. Although it is unclear which DRG neuron subtypes are infected during natural VZV infection, it is highly likely that nociceptive DRG neurons are susceptible to VZV infection and host a latent state. This supports and extends previous observations found in the SCID-hu DRG mouse model^46^. VZV-infected mHD10.6 neurons produced infectious virus particles that disseminated through the culture via cell-to-cell spread. However, similar to other neuronal models of VZV infection^17,47,48^, we also observed minimal CPE in VZV-infected mHD10.6 neurons. These observations contrast with lytic VZV infection in fibroblast, epithelial, or melanoma cells, where VZV infection inevitably results in cell rounding cytopathic effects, and in some cell types, extensive syncytium formation and destruction of the infected cells within days^49,50^. In addition to differential susceptibility to apoptosis^51^, the delayed and reduced replication of VZV in mHD10.6 neurons compared to other cell types could explain these observations.

VZV latency can be defined by the maintenance of the viral genome in host cells in the absence of virus production, while maintaining the capacity to reinitiate lytic VZV infection. The quiescent-persistent VZV latent state in mHD10.6 neurons recapitulates essential features of VZV latency in human TG. VZV genomes are maintained at stable levels over a period of at least 20 days, are transcriptionally repressed, and no infectious virus could be recovered from the cultures. While expression of all VZV genes was reduced, the pattern of viral gene expression changed with prolonged VZV latency (20 dpi), where transcripts originating from the VLT–ORF63 region were less reduced compared to the analyzed lytic VZV genes. Possibly, the selective expression of *VLT* and *VLT-ORF63* observed *in vivo* follows a more gradual repression of VZV gene expression, requiring a longer culture of latently-infected mHD10.6 cells. It is not clear whether the continuous low-level transcription of lytic VZV genes occurs in all cells harboring latent viral genomes, or if it reflects occasional episodes of transcriptional derepression reminiscent of HSV phase I reactivation^52^. The observation that in some wells one or a few neurons continued to express lytic proteins and GFP indicates heterogeneity in VZV DNA silencing between infected cells, possibly due to variability in viral genome chromatinization^10,11^. However, such cultures never produce spreading virus, even when cocultured with highly VZV susceptible ARPE19 cells.

Previous studies in other VZV model latent states have also shown the maintenance of genomes following infection in the presence of antivirals to prevent viral DNA replication and repress viral gene expression^17,18,20,23,33^. Here, we tested the nucleoside analogs ACV, BVDU, and the more recently developed helicase-primase complex inhibitor AMNV^41–43,53,54^. All compounds were able to induce quiescent persistent VZV infection, although ACV and BVDU more potently inhibited lytic gene expression. However, additional research is needed to determine if the choice of antiviral impacts the efficiency of VZV reactivation. ACV and BVDU are potentially incorporated into replicating VZV DNA^34,34–36,53–55^, which could lead to replication intermediates and latent genomes carrying these nucleoside analogs. By contrast, AMNV prevents the initiation of VZV DNA replication and is expected to leave the viral genome intact^41–43,53,54^.

We obtained a small fraction of latently infected cultures showing a reactivation event reproducibly using capsaicin treatment, in line with the expression of the receptor in the mHD10.6 cells^24^. Intriguingly, not all treatments resulted in stimulation of reactivation events, including some that have been shown to reactivate latent VZV in other model neuronal platforms. The triggers and mechanisms involved in VZV reactivation in humans remain unclear. Previous studies showed that VZV reactivation can be induced by inhibition of nerve growth factor signaling, activation of c-Jun N-terminal kinase (JNK) signaling via PI3K-inhibition, or inhibition of histone deacetylases in cultures of stem cell-derived neurons^17–19^. Remarkably, none of these stimuli was able to efficiently induce VZV reactivation in the mHD10.6 neurons. Based on the original publication, the HD10.6 neurons revealed capsaicin sensitivity and excitability, where the addition of capsaicin to the culture medium induced cobalt uptake and electrophysiological changes^24^, we found that the addition of capsaicin to the culture medium was the only reproducible compound that triggered reactivation of VZV. This data is in line with previously reported findings where neuron excitability has been shown to influence HSV-1 reactivation^56,57^. Due to the broad effects of the increased excitability of the neurons, it is difficult to speculate on the exact mechanisms by which capsaicin induces reactivation.

Despite finding stimulation, or reactivation events, it was notable that only a fraction of the latently VZV-infected neuron cultures responded to capsaicin to reactivate and spread throughout the culture. The inefficient low level of reactivation was also reported in other human neuron model systems^17,18,23^. However, we note that this *in vitro* reactivation is not unlike *in vivo* reactivation of VZV in humans, where clinical reactivation of VZV occurs only in a third of VZV-infected individuals and mostly occurs only once in a person’s lifetime, and likely originates from one single reactivated cell^58,59^. The lack of reactivation in a multitude of neurons has also been described in the hESC, iPSC, HSN models, and SH-SY5Y cell line for VZV, and also in HSV-1^17,18,20,23,60,61^. This suggests that VZV latency is a highly efficient process, which results in a reduced frequency of VZV reactivation compared to other alphaherpesviruses.

In conclusion, our study shows that the mHD10.6 model provides an easy-to-use, clonal, and scalable human sensory neuron model to study lytic, latent-like, and reactivated VZV infection. The mHD10.6 model facilitates functional studies on VZV neuronal infection and holds potential to allow manipulation for more in-depth research regarding virus and host targets of therapeutic value to limit the establishment of VZV latency and potentially to prevent reactivation.

## Materials and Methods

### Cell culture

HD10.6 cells were a kind gift from Dr. Anna Cliffe (University of Virginia) and maintained in 75 cm^2^ cell culture flasks (Corning), precoated for 4 hours at 37 °C with 16.67 µg/mL fibronectin (Merck) in PBS. Cells were grown in proliferation medium, composed of Advanced Dulbecco’s modified eagle medium/Ham’s F12 (DMEM/F-12; Gibco) containing 1x GlutaMAX (LIFE), 1x B-27 (LIFE), 10 ng/mL prostaglandin E1 (Merck), 100 µg/mL Primocin (Invivogen), and 0.5 ng/mL recombinant human fibroblast growth factor basic (FGF-b; Peprotech). For maturation, HD10.6 cells were seeded at 2.5×10^4^ cells/cm^2^ in proliferation medium on plates precoated with 2% (v/v) Matrigel (Corning) and 10 µg/mL poly-D-lysine (Merck) in PBS (day 0). At day 1, medium was replaced by maturation medium, composed of neurobasal A medium (Gibco) supplemented with 1x GlutaMAX, 1x NCS21 (Capricorn), 1 µg/mL doxycycline (Merck), 50 ng/mL nerve growth factor (NGF; Peprotech), 25 ng/mL ciliary neurotrophic factor (CNTF; Peprotech), 25 ng/mL glial cell-derived neurotrophic factor (GDNF; R&D systems), and neurotrophin-3 (NT-3; Peprotech). Every 3-4 days, 50% of the medium was replaced by fresh maturation medium. Retinal pigmented epithelial cells (ARPE-19) were grown in a 1:1 (v/v) mix of DMEM and Ham’s F12 containing 10% (v/v) heat-inactivated fetal bovine serum (FBS; Merck), 1x penicillin/streptomycin (Capricorn), and 2 mM L-glutamine (Capricorn). All cells were incubated at 37 °C and 5% (v/v) CO_2_. HD10.6 and ARPE-19 cells were confirmed to be negative for mycoplasma and authenticated by short tandem repeat profiling **(Supplementary Table 1)**.

### Generation of VZV rpOka.ORF49.T2A.GFP (VZV-49GFP)

Because the original pOkaBAC clone contained several non-synonymous substitutions^62^, we generated a new self-excisable pOkaCre clone that contained the virulent pOka 2096 G residue in ORF31, but not any of the non-synonymous substitutions (**Supplementary Materials and Methods**). Stellar *E. coli* harboring the pOkaBACre genome were transformed with the pST76A-SR shuttle plasmid containing the ORF49-T2A-EGFP cassette (pST76A-SR_ORF49T2AGFP, **Supplementary Materials and Methods**), and recA-mediated allelic exchange was carried out as described previously^63^, resulting in pOkaBACreORF49T2AGFP/Stellar. The pOkaBACreORF49T2AGFP genome was extracted (Genopure Plasmid Maxi Kit, Roche), subjected to restriction fragment length polymorphism analysis using BamHI or EcoRI in parallel with the original pOkaBAC and pOkaBACre genomes, and the region used for allelic exchange was sequenced. The purified pOkaBACreORF49T2AGFP genome (1 µg) was transfected into ARPE-19 cells (1 x 10^5^) using PEImax solution (3 µL) prepared as described previously^12^. At 6 days post-transfection, a typical cytopathic effect with GFP expression driven by ORF49 promoter was seen, establishing recombinant pOkaORF49T2AGFP virus (VZV.rpOka.ORF49.T2A.GFP), then cell-free virus was prepared as described previously^29,64^. Self-excision of “BACre” cassette flanked by loxP by endogenous Cre recombinase was confirmed by PCR of genomic DNA extracted from rpOkaORF49T2AGFP-infected ARPE-19 cells using primers ORF11F1794 and ORF12R211 (**Table 2**).

**Table 2.**
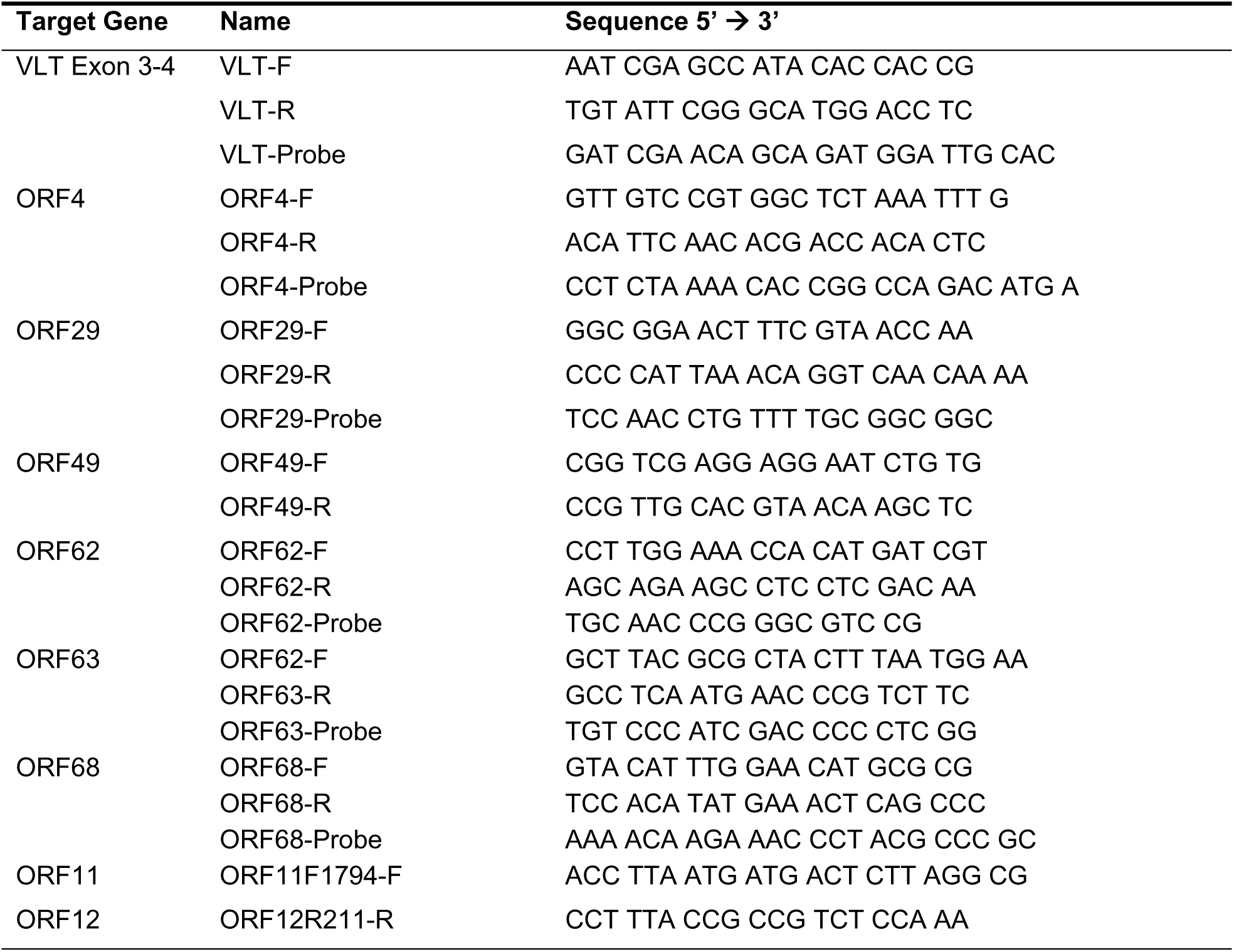
List of used primers.

### Generation of VZV POka-57GFPLZ

VZV pOka-57GFPLZ was used in some latency and reactivation studies, and its construction will be detailed separately in a manuscript (Flot and Kinchington, in preparation). Briefly, a cell line was created based on ARPE-19 cells that express Strep pyogenes CRISPR-Cas9 and guide RNAs to target the intergenic sequences of ORF56 and ORF57. VZV POka infected ARPE19-Cas9sgRNA cells were transfected with an ORF57GFP construct with additional flanking sequences to allow recombination into the VZV genome. Resultant fluorescent plaques were plaque-purified to purity and a homogeneous population.

### VZV infection of mHD10.6 neurons

Cell-free virus was produced from VZV-49GFP-infected ARPE-19 cells showing about 50% CPE, and VZV-57LZ-GFP from infected htRPE cells as previously described^65^. Fresh cell-free virus stocks were made by scraping the cells in base maturation medium (Neurobasal A medium, supplemented with 1x GlutaMAX), followed by sonication and clarification for 15 minutes at 1000xg for 15 minutes. VZV-49GFP virus stocks were immediately used for infection of HD10.6 cells and titration on ARPE-19 cells. VZV-57LZ-GFP stocks were concentrated using 1:4 Lenti-X™ Concentrator (TaKaRa) and resuspended in 1:10 initial volume PGSC, before quick freeze in liquid nitrogen, and subsequently frozen at -80 °C. When needed, VZV57LZ-GFP stocks were rapidly thawed, diluted to the appropriate concentration in neuronal differentiation media, and immediately used for infection on HD10.6 cells for four hours, followed by washing of cells and media replenishment. For lytic infection, mHD10.6 were infected with 500-1000 PFU/well in a 24-well format for two hours in base maturation medium, after which the inoculum was removed and replaced by complete maturation medium. For latent infections, both the infection and culture of the VZV-infected HD10.6 were performed in the presence of 300 µM acyclovir (ACV; Merck), 2 µM BVDU (Brunschwig Chemie BV), or 1 µM AMNV (MedChem Express). Every 2-3 days, half-volume medium changes were performed by adding complete differentiation medium containing 300 µM ACV for a total of five or ten days, after which cells were cultured in complete maturation medium without antivirals for 7-10 more days.

### Reactivation of latent VZV

To establish latent infection for reactivation, mHD10.6 were infected 1000 PFU/well on 24-well plates initially seeded with 25,000 cells for four hours at 33°C with 5% CO2 in complete maturation media, in the presence of 300 µM ACV, 2 µM BVDU, and 1 µM AMNV (all from MedChem Express). Following the 4-hour infection period, virus-containing media were removed, and cultures were gently washed once with 1X PBS. The cultures were then maintained with complete maturation media with antivirals for ten days. The antiviral withdrawal period was initiated by replacing antiviral-containing media with 1x PBS and adding complete maturation media. Media was replaced by 50% replenishment every two days for a period of 7 to 10 days. Neurons were monitored every few days using whole plate automated imaging scans using an Olympus inverted microscope live cell microscope (Olympus IX80 Inverted Microscope, CellSens software). To reactivate latently infected mHD10.6 cells were treated with media containing 20 µM LY294002 (48h; MedChem Express), medium lacking NGF (48h), 5 mM sodium butyrate (168h; Sigma-Aldrich), 5 µM JQ-1 (48h; MedChem Express), 60 µM forskolin (48h; TOCRIS), medium lacking doxycycline (Sigma-Aldrich), 10 µ g/mL ADU-S100 (48h; InvivoGen), 30 µM capsaicin (48h; MedChem Express), (20 µM) 12-O-Tetradecanoyl-phorbol-13-acetate (TPA; 168h; MedChem Express), 43 °C heat shock (3h), or VZV superinfection (MOI 0.01; 240h) in complete media (or other stimulus as detailed in the text or **Figure 1**) followed by media changes at 48hrs post induction with maturation media.

### Virus titration

The titer of VZV-49GFP stocks was determined by performing a 4-fold serial dilution, in 8 technical replicates per plate with 2 plates, on monolayers of ARPE-19 cells in 96-well cell culture plates. GFP-positive wells were scored using a fluorescence microscope, and the titer was calculated using the Spearman-Karber method^66,67^.

### Transmission electron microscopy

Mock- and VZV-infected mHD10.6 cells cultured in 6-cm dishes were fixed for an hour at room temperature by adding double-concentrated fixative to the medium (final concentration: 1.5% glutaraldehyde in 0.1M cacodylate buffer). After 3x rinsing with 0.1 M cacodylate buffer, the cells were postfixed with 1% osmium tetroxide and 1% uranyl acetate. Cells were dehydrated with a series of ethanol, followed by a series of mixtures of ethanol and EPON (LX112), and at the end pure EPON. BEEM capsules filled with EPON were placed on the dishes with the open face down. After EPON polymerization at 45°C for the first night and 70°C for the second night, the BEEM capsules were snapped off. Ultrathin sections 100 nm were made parallel to the surface of the BEEM capsules containing the cultured cells using a Leica EM Ultracut 6. The sections were contrasted with 7% uranylacetate (20min) and Reynolds lead citrate (10min) and examined at 120 keV with a Tecnai Twin transmission electron microscope (Thermo Fisher). Overlapping images were automatically collected with a Gatan Oneview camera (Gatan) at binning 2 at a nominal magnification of 11000 x (1.96 nm pixel size) and stitched together into a composite image as previously described^68^.

### Immunofluorescence

HD10.6 were cultured on 8-well microscope chamber slides for immunofluorescence. mHD10.6 were fixed using 4% (w/v) paraformaldehyde (PFA; Santa Cruz Biotechnology) for 15 minutes at 4 °C, permeabilized in 0.1% (v/v) Triton X-100 (Merck) in PBS for 30 minutes at room temperature (RT) and blocked using 5% (v/v) donkey serum (Merck) for 30 minutes at 4 °C. Cells were then incubated with 1 µg/mL rabbit-anti-beta-III-tubulin (BioLegend), 4 µg/mL goat-anti-TrkB (BioTechne), and 1:250 mouse-anti-peripherin (Merck), or 1:250 rabbit-anti-cleaved caspase-3 (Cell Signaling Technologies) for 1 hour at 4 °C in the dark. Cells were washed three times using PBS, followed by incubation with 1:500 donkey anti-goat AlexaFluor-488 (Invitrogen), 1:500 donkey anti-mouse AlexaFluor-555 (Invitrogen), and 1:500 chicken anti-rabbit AlexaFluor-647 (Invitrogen) for 1 hour at 4 °C in the dark, and 5 µg/mL Hoechst 33342 (LIFE) was added for the last 15 minutes. Fluorescent images were made using a confocal microscope (Carl-Zeiss LSM 770) and analyzed using Zen blue version 3.4 (Carl-Zeiss).

### RNA isolation, cDNA synthesis, DNA isolation, and (RT-)qPCR

HD10.6 cells were cultured in 24-well plates. RNA isolation was performed using the E.Z.N.A HP total RNA kit (Omega Bio-Tek) following the manufacturer’s protocol. RNA samples were treated using TURBO DNA-free (Ambion) according to the manufacturer’s protocol. cDNA synthesis was performed using SuperScript IV (LIFE) with random primers (Promega), dNTPs (Merck), and RNAsin (Promega). Synthesized cDNA was diluted 10-fold in RNAse-free DEPC ultrapure H_2_O (LIFE). DNA was isolated using E.Z.N.A. Tissue DNA Kit (Omega Bio-Tek) following the manufacturer’s protocol. (RT-)qPCR was performed using 4x TFA mix with specific primer/probe mixes against specific primers *TUBB3* (IDT; Hs.PT.58.20385221)*, NTRK2* (IDT; Hs.PT.58.41039984)*, PRPH* (IDT; Hs.PT.58.25317355)*, ACTB* (IDT; Hs.PT.39a.22214847)*, RET (IDT;* Hs.PT.58.4318912) and primers found in **Table 1** (*VLT, ORF63, ORF68, ORF4, ORF29, ORF62, ORF49, VZV-Termini*). For reactivation experiments, DNA and RNA were isolated using Zymo Research Quick-DNA/RNA Miniprep Kits (Zymo) following the provided protocol for extraction from cells. The cDNA for these experiments was synthesized using the iScript™ cDNA Synthesis Kit (BIORAD). Synthesized cDNA resuspended in nuclease-free H_2_O. RT-qPCR and qPCR were performed using the SsoAdvanced Universal SYBR Green Supermix (BIORAD).

### Statistical analysis

All analyses were performed in GraphPad Prism 10.2.2 (397) (Graphpad Software LLC, California, USA). All statistical analyses were done by grouped data 2-way ANOVA using the Geisser-Greenhouse correction model and corrected for multiple comparisons using Tukey’s statistical hypothesis testing.

## Acknowledgements

The authors would like to thank Dr. Anna R. Cliffe (University of Virginia, Charlottesville, VA, USA) for sharing the HD10.6 cells and protocols. ACH, GMGMV, PRK, and WJDO are supported by the National Institute of Allergy and Infectious Diseases (NIAID) of the National Institutes of Health (NIH) under Award Number R01-AI151290. PRK acknowledges support for this work by R01 AI158510; P30 EY08098; The Eye & Ear Foundation of Pittsburgh, and the Research to Prevent Blindness Inc. NY. TS is supported by the Ministry of Education, Culture, Sports, Science and Technology (MEXT KAKENHI JP21H02741 and JP23K21379) and SHIONOGI INFECTIOUS DISEASE RESEARCH PROMOTION FOUNDATION (a grant for fundamental research, 2024F004).

